# Oxytocin has sex-specific effects on social behaviour and hypothalamic oxytocin immunoreactive cells but not hippocampal neurogenesis in adult rats

**DOI:** 10.1101/859165

**Authors:** Paula Duarte-Guterman, Stephanie E. Lieblich, Wansu Qiu, Jared E.J. Splinter, Kimberly A. Go, Laura Casanueva-Reimon, Liisa A.M. Galea

**Affiliations:** Djavad Mowafaghian Centre for Brain Health, University of British Columbia, Vancouver, BC, Canada; Department of Psychology, University of British Columbia, Vancouver, BC, Canada; Graduate Program in Neuroscience, University of British Columbia, Vancouver, BC, Canada

**Author notes:** Address all correspondence and requests for reprints to: L. A. M. Galea, PhD, Djavad Mowafaghian Centre for Brain Health, 2215 Wesbrook Mall, Vancouver, British Columbia V6T 1Z3, Canada,.

**Keywords:** social investigation, sex differences, dentate gyrus, hypothalamus, nanoparticle, estradiol, females, hippocampus, doublecortin

## Abstract

Oxytocin regulates social behaviours, pair bonding and hippocampal neurogenesis but most studies have used adult males. Our study investigated the effects of oxytocin on social investigation and adult hippocampal neurogenesis in male and female rats. Oxytocin has poor penetration of the blood-brain barrier, therefore we tested a nanoparticle drug, TRIOZAN^TM^ (Ovensa Inc.), which permits greater blood-brain-barrier penetration. Adult male and female rats were injected daily (i.p.) for 10 days with either: oxytocin in PBS (0.5 or 1.0 mg/kg), oxytocin in TRIOZAN^TM^ (0.5 or 1.0 mg/kg), or vehicle (PBS) and tested for social investigation. Oxytocin decreased body mass and increased social investigation and number of oxytocin-immunoreactive cells in the supraoptic nucleus (SON) of the hypothalamus in male rats only. In both sexes, oxytocin decreased the number of immature neurons (doublecortin+ cells) in the ventral hippocampus and reduced plasma 17β-estradiol levels in a dose- and delivery-dependent way. Oxytocin in TRIOZAN^TM^ reduced sedation observed post-injection and increased some central effects (oxytocin levels in the hypothalamus and ventral hippocampus neurogenesis) relative to oxytocin in PBS indicating that the nanoparticle may be used as an alternative brain delivery system. We showed that oxytocin has sex-specific effects on social investigation, body mass, sedation, and the oxytocin system. In contrast, similar effects were observed in both sexes in neurogenesis and plasma 17β-estradiol. Our work suggests that sex differences in oxytocin regulation of brain endpoints is region-specific (hypothalamus versus hippocampus) and that oxytocin does not promote social investigation in females.

## Introduction

Oxytocin is well known for regulating social behaviours and pair bonding (Lim and Young, 2006) and is mainly synthesized and secreted in the hypothalamic paraventricular nucleus (PVN) and supraoptic nucleus (SON). Surprisingly, few studies have examined roles for oxytocin in both sexes even though there are sex differences in the effects of oxytocin on social behaviour and brain activation in both humans and animal models (DiBenedictis et al., 2017; Dumais et al., 2017, 2016b; Lukas and Neumann, 2014; Rilling et al., 2014; Smith et al., 2017; Steinman et al., 2016; and reviewed by Bredewold and Veenema, 2018; Caldwell, 2018). Social behaviour regulation is associated with a network that includes the prefrontal cortex, medial amygdala, bed nucleus of the stria terminalis and hypothalamus (Ko, 2017) and many of these areas are also connected to the hippocampus (reviewed in Jurek and Neumann, 2018). Thus, perhaps it is not surprising that the integrity of the hippocampus has been implicated in social cognition (Alexander et al., 2016; Danjo et al., 2018; Kumaran et al., 2016; Matta et al., 2017; Okuyama et al., 2016; Omer et al., 2018; Raam et al., 2017). In mammals, the hippocampus retains the ability to produce new neurons throughout the lifespan (Kempermann et al., 2018). New neurons in the hippocampus have been implicated in pattern separation, stress resilience, and anxiety-related behaviours (Anacker et al., 2018; Clelland et al., 2009; Snyder et al., 2011). Furthermore, social isolation (Leasure and Decker, 2009; Stranahan et al., 2006), social stress, (McCormick et al., 2012, 2010) and social interactions (Peragine et al., 2016, 2014; Tzeng et al., 2016) alter neurogenesis in the hippocampus. Hormones, such as sex steroids and prolactin, regulate different aspects of neurogenesis, proliferation, differentiation and survival of new neurons, in a sex-dependent way (reviewed in Mahmoud et al., 2016). In male rodents, oxytocin promotes social behaviour (Lukas et al., 2011; Ramos et al., 2013) and increases adult hippocampal neurogenesis (Leuner et al., 2012; Opendak et al., 2016; Sánchez-Vidaña et al., 2016). However, it is unknown whether oxytocin regulates hippocampal neurogenesis in females. Oxytocin receptors (OTR) are expressed in the hippocampus of both sexes in rodents (Dumais et al., 2013; Lin et al., 2017; Yoshida et al., 2009). However, there are sex differences with female rats having lower OTR binding densities in many regions including the CA1 of the hippocampus compared to males (Dumais et al., 2013). Furthermore, oxytocin activates brain regions differently in males and females (Dumais et al., 2017) and it is possible that the regulation of hippocampal neurogenesis and social investigation by oxytocin is different between the sexes.

Oxytocin is a large hydrophilic peptide, which has poor penetration of the blood-brain barrier (BBB) and a very short half-life in the blood (∼3-9 min) (Churchland and Winkielman, 2012; Leng and Ludwig, 2016; Modi and Young, 2012). Following systemic oxytocin, only up to 1.3% of peripheral oxytocin enters the brain (Ermisch et al., 1985; Mens et al., 1983). The use of a nanotechnology drug delivery system that both permits BBB penetration and lengthens the half-life may offer greater delivery of oxytocin to the brain. TRIOZAN^TM^ nanomedicine delivery platform (Ovensa Inc) is based on a high value-added chitosan derivative called trimethyl chitosan (TMC), which enables efficient encapsulation resulting in protection of drug molecules against degradation, effectively maintaining therapeutic integrity (Kulkarni et al., 2017; Mourya and Inamdar, 2008; Ovensa Inc., 2018). TRIOZAN^TM^ is a safe, biodegradable and biocompatible biopolymer and has the ability to accumulate in the targeted tissues with minimal loss, enabling controlled release of therapeutics via effective penetration of the BBB (Ovensa Inc., 2018). TMC nanoparticles have been previously used for drug delivery to different tissues, including the brain (Cardia et al., 2019; Kumar et al., 2013; Wang et al., 2010), because TMC enhances transport of large molecules through the adsorptive-mediated transcytosis mechanism (Lu, 2012; Ovensa Inc., 2018). Therefore, in the present study we compared the effects of systemic oxytocin with and without the nanoparticle TRIOZAN^TM^.

The objective of this study was to investigate whether systemic oxytocin delivered in vehicle or with a nanoparticle drug delivery platform (TRIOZAN^TM^) regulates social investigation, oxytocin levels in the brain and periphery, oxytocin-immunoreactive cells in the hypothalamus (SON and PVN) and adult hippocampal neurogenesis in male and female rats. We hypothesized that oxytocin would impact the sexes differentially and that TRIOZAN^TM^ would increase delivery of oxytocin to the brain and result in enhanced effects on social investigation and hippocampal neurogenesis.

## Materials and Methods

### Animals and treatments

Adult male and female Sprague–Dawley rats (n=5-8/treatment/sex; 60 days old and tested around 80 days old) were obtained from Charles River (Quebec, Canada) and double-housed with food and water available ad libitum. All protocols were approved by the Institutional Animal Care Committee at the University of British Columbia and conformed to the guidelines set out by the Canadian Council for Animal Care. Ten days after arrival, males and females were randomly divided into one of the following treatment groups: oxytocin in 0.1 M PBS (0.5 or 1.0 mg/kg), oxytocin delivered in TRIOZAN^TM^ (0.5 or 1.0 mg/kg, prepared by PO-Laboratories, Dorval, QC, Canada), and vehicle control (equivalent volume of 0.1 M PBS, pH= 7.4). The volume of injection was 1.0-1.2 ml for females and 1.5-2.0 ml for males. Oxytocin encapsulated in TRIOZAN^TM^ was prepared using PBS and the resulting solution contained both free (dissolved) and encapsulated oxytocin. Concentrations of free and encapsulated oxytocin were determined by high performance liquid chromatography (HPLC) and resulted in 42.2% of oxytocin encapsulated in TRIOZAN^TM^ (PO-Laboratories). Animals received daily i.p. injections for 10 days between 9:00 and 11:00. As oxytocin injections can cause sedation (e.g., Uvnäs-Moberg et al., 1994; the current study), we monitored sedation 5-10 mins post-injection over the course of the 10 days using a binary score (0 = no sedation, 1 = sedation indicated by a clear lack of locomotor activity and rearing, eyes closed and rats appearing asleep). In addition, animals were behaviourally tested two hours after injection on day 9 to avoid confounding sedative and/or muscular effects of oxytocin that would affect locomotion (Elabd et al., 2014; Hess et al., 2016; Leong et al., 2016; Uvnäs-Moberg et al., 1994). On day 10, brains were collected 30 mins after oxytocin injection. This timeline coincides with peak levels in microdialysates sampled within amygdala and hippocampus (after an i.p. injection of oxytocin in male mice; Neumann et al., 2013). Similar nanoparticles to TRIOZAN^TM^, such as TMC, increase delivery of drugs within 30 mins (after inhalation of progesterone bound to TMC in male mice; Cardia et al., 2019) and between 60-240 mins (after injection with anti-neuroexcitation peptide bound to TMC in male mice; Wang et al., 2010). Therefore, we chose 30 mins as it was within the range of previous studies using a nanoparticle to deliver a drug to the brain and the temporal profile of oxytocin in saline.

### Social Investigation

The three-chambered social paradigm allows for the measurement of social investigation in rodents (Crawley, 2004; Ku et al., 2016; Nadler et al., 2004). On day 9, two hours after injection, animals were allowed to explore a novel arena (100 cm x 40 cm x 40 cm) composed of three connecting chambers separated by open sliding doors (Figure 3) for 10 min. Locomotion, grooming, rearing, behaviours were assessed during this time to evaluate general activity. Time spent in each of the chambers was also measured to ensure animals did not have a preference for a particular side of the chamber. Following this habituation period, animals were briefly kept (for ∼2-3 min) in the central chamber with sliding doors closed while an unfamiliar and novel same sex conspecific rat was placed in one of the outer chambers under a small wire cage. An identical empty wire cage was secured in the other outer chamber. The unfamiliar rat was randomly assigned to either the right or left side of the chamber. The test animal was then allowed to explore the three chambers for a further 10 min. All sessions were recorded and scored by an observer unaware of treatment conditions. Videos were scored for (1) time spent in the chamber that housed the novel conspecific rat (i.e., social stimulus rat) and (2) time spent engaging in investigatory behaviour with the novel rat (sniffing and touching the basket where the novel rat was confined; i.e. social investigation).

### Tissue and blood collection

Vaginal lavages were conducted on days 9 (before behaviour) and 10 (at euthanasia) to account for estrous cycle phase effects on our measures. On day 10, 30 mins after receiving the treatment, all rats received a lethal overdose of sodium pentobarbital. Blood was collected via cardiac puncture into cold EDTA coated tubes and centrifuged 4 h later for 10 min at 4°C. Plasma was stored at −80°C until ELISA and radioimmunoassay (RIA). Adrenals were extracted and weighed. Hypothalamus was quickly dissected from the left hemisphere, flash frozen on dry ice, and stored at −80°C. The right hemisphere was fixed for 24 h in 4% paraformaldehyde in 0.1M PBS (4°C) and then transferred to a 30% sucrose/0.1 M phosphate buffer solution for cryoprotection until brains sunk. Serial coronal sections (30 μm) were cut with a freezing microtome (SM2000R; Leica, Richmond Hill, ON) across the extent of the hippocampus (collected in 10 series) and stored in an antifreeze solution (containing 0.1M PBS, 30% ethylene glycol, and 20% glycerol) and stored at −20°C.

### Oxytocin and 17β-estradiol concentrations

Oxytocin concentrations were analyzed using an enzyme-linked immunosorbent assay kit (Enzo Life Sciences, ADI-900-153A) on plasma and hypothalamus homogenate samples. Plasma samples were run in duplicate according to the manufacturer’s instructions. Hypothalamus (and other brain regions, frontal region and hippocampus) samples were homogenized in 200μl of cold Tris lysis buffer (150 mM NaCl, 20 mM Tris, pH=7.5, 1 mM EDTA, 1 mM EGTA, 1% Triton X) containing a cocktail of protease and phosphatase inhibitors using a bead disruptor (Omni, VWR). Homogenates were centrifuged at 2100g for 10 min at 4°C and supernatants stored at −80°C. Hypothalamus homogenates were run in duplicate on a modified ELISA using the lysis buffer to prepare the standard curve (Enzo Life Sciences). Oxytocin levels were normalized to protein content using a BCA protein assay (ThermoFisher) and expressed as pg/mg protein. The oxytocin antibody is highly specific with 7.5% cross-reactivity with Arginine-Vasotocin. The sensitivity of the assay is 15 pg/ml and average intra- and inter-assay coefficients of variation were <10%. 17β-estradiol levels were determined in plasma using a commercially available radioimmunoassay kit (ultrasensitive estradiol RIA, Beckman Coulter, DSL4800). All samples were run in duplicate. The assay sensitivity is 2.2 pg/ml and average intra-assay coefficient of variation was <20%. Samples below the zero standard curve point were replaced with 0 pg/ml.

### Doublecortin Immunohistochemistry

Free floating sections were rinsed three times for ten minutes in 0.1M phosphate buffered saline (PBS; pH=7.4) in between each step listed below, unless otherwise stated. Endogenous peroxidase activity was quenched with a 30 min incubation in 0.6% H_2_O_2_. Tissue was incubated in a goat anti-DCX polyclonal antibody (1:1000 in PBS containing 3% normal rabbit serum and 0.4% Triton X-100; Santa Cruz Biotechnology, Cat# sc-8066, RRID: AB_2088494) for 24 h at 4°C. Tissue was rinsed five times for ten minutes in PBS and then incubated overnight at 4°C in a rabbit anti-goat secondary (1:500; Vector Laboratories) diluted in PBS. Sections were then washed five times for ten minutes in PBS and then incubated in an avidin-biotin horseradish peroxidase complex (1:1000; ABC Vector Elite kit; Vector Laboratories) for 4 h at room temperature. Tissue was then washed in sodium acetate buffer (0.175 M; pH= 7.3) two times for 2 min each before reacting with diaminobenzidine in the presence of nickel (DAB Peroxidase Substrate Kit, Vector Laboratories) according to the manufacturer instructions. Sections were mounted onto Superfrost/Plus slides (Fisher Scientific, Ontario, Canada), dried several days, dehydrated in ethanol and cleared with xylene, and then cover slipped with Permount (Fisher Scientific). The primary antibody was omitted in a control sample and did not contain any immunoreactivity.

### Oxytocin immunofluorescence

Free floating sections were rinsed three times for ten minutes in 0.1M phosphate buffered saline (PBS; pH 7.4) in between each step listed below, unless otherwise stated. Sections were incubated for 10 min in 0.1% sodium borohydride in PBS on ice and then blocked in PBS containing 10% normal goat serum (NGS) for 30 min. Sections were then incubated in mouse anti-oxytocin monoclonal antibody (1:2000, Millipore, Cat# MAB5296, RRID: AB_2157626) in 3% NGS and 0.5% Triton-X for 44 h at 4°C and then in goat anti-mouse secondary antibody Alexa 488 (1:500; Invitrogen) in 3% NGS and 0.5% Triton-X for 2 h. After four rinses in PBS, sections were mounted onto Superfrost/Plus slides (Fisher Scientific) and coverslipped using polyvinyl alcohol with DABCO (Sigma-Aldrich).

### Microscopy

A researcher blinded to experimental conditions counted doublecortin (DCX)+ cells using a standard light microscope. DCX+ cells were counted in the entire granule cell layer of the right hippocampus at 400x magnification. Sections were considered ventral when the dentate gyrus at the bottom of the section was obviously present, that is between −5.2 mm to −6.7 mm Bregma (Paxinos and Watson, 2005). DCX is a microtubule-associated protein expressed in proliferating progenitor cells and immature neurons, from 2 h to 3 weeks after birth in rats (Brown et al., 2003). DCX+ cells can be classified into developmental stages: proliferative (type 1, no or short processes present), intermediate (type 2, medium process with no branching), or post-mitotic (type 3, branching dendrites; Figure 4A). For DCX+ cell phenotyping, a total of 120 cells per rat were randomly selected from three different sections from dorsal (60) and ventral (60) regions of the hippocampus (Gobinath et al., 2016; Workman et al., 2016). Total GCL volumes (mm^3^) from dorsal and ventral sections were quantified from digitized images using ImageJ (U. S. National Institutes of Health, Bethesda, Maryland, USA, http://imagej.nih.gov/ij/). OT-immunoreactive (ir) cells were counted at 400x magnification in one section of the supraoptic nucleus (SON) and one section of the paraventricular nucleus (PVN) of the hypothalamus for each brain using an epifluorescence microscope. Sections of the SON (around Bregma −1.44 and −1.56 mm) and PVN (around Bregma −1.80 and −1.92 mm; Paxinos and Watson, 2005) were anatomically matched across all animals.

### Statistical Analysis

All analyses were performed using Statistica v.8.0 (StatSoft Inc, Tulsa, OK). Time spent in the social stimulus chamber (time spent on side of the chamber containing novel rat divided by total time in the arena), time spent investigating the stimulus rat (time spent sniffing and touching the basket containing novel rat divided by total time in the arena), and maturity of DCX+ cells were transformed before analysis using the arcsine transformation. The oxytocin in plasma and hypothalamus data were log transformed and the 17β-estradiol data were square root transformed to meet normality and heteroscedasticity requirements. All data (unless specified) were analyzed using ANOVA with sex (male, female) and treatment (PBS control, oxytocin in PBS (0.5 or 1.0 mg/kg), oxytocin in TRIOZAN^TM^ (0.5 or 1.0 mg/kg)) as between-subject factors. Number (number of cells divided by number of sections counted) and maturity of DCX+ cells, and GCL volume were each analyzed using repeated measures ANOVA with hippocampal region (dorsal, ventral) and an additional within-subjects factor for maturity of DCX+ cells (proliferative, intermediate, post-mitotic). The data for OT-ir cells in the SON were not normal or homoscedastic with female counts being more variable than male counts, even after transformation. We therefore analyzed males and females separately to meet ANOVA assumptions. 17β-estradiol data were analyzed using ANOVA separately for both sexes and using estrous cycle as between-subject factors in females. We identified estrous cycle phase but due to low sample sizes (N=1-2 for some treatment groups in each phase) we used estrous cycle stage or 17β-estradiol as a covariate on our measures and the covariate effect is only presented when significant. When appropriate, post-hoc analysis used the Neuman-Keul’s procedure. Test statistics were considered significant if p ≤ 0.05. A priori comparisons were subject to Bonferroni corrections. Pearson’s correlations were performed between variables of interest.

## Results

### Body and adrenal mass

Body mass immediately prior to the start of the injections did not differ between treatment groups (all p’s>0.3) however males (405.6g ± 4.8g (SEM)) were larger than females (mean±SEM= 252.5±1.6 g; Cohen’s *d* = 7.31; main effect of sex F(1,58)=829.93; p<0.0001; partial η^2^= 0.93; Table 1). Net body mass gain (last day minus first day of injections) was reduced in rats treated with oxytocin in PBS relative to the PBS controls (0.5 and 1.0 mg/kg doses; p=0.04 and 0.003, respectively; main effect of treatment (F(4,58)=6.16; p<0.001; partial η^2^= 0.30) but this effect was driven by the males. In males, oxytocin in PBS or in TRIOZAN^TM^ (0.5 and 1.0 mg/kg) reduced body mass gain relative to the PBS male controls (a priori comparisons, oxytocin in PBS by dose: p=0.01 and p=0.0003; Cohen’s *d* = 1.15 and 1.46; oxytocin in TRIOZAN^TM^ by dose: p’s<0.0001; Cohen’s *d* = 2.04 and 1.88, respectively), while oxytocin treated females were not affected relative to PBS female controls (p’s>0.14, respectively). No significant differences were observed between oxytocin in PBS and oxytocin delivered in TRIOZAN^TM^ (p’s>0.2). There was also a sex difference with males having a larger body mass gain than females (main effect of sex F(1,58)=32.14; p<0.0001; partial η^2^ = 0.36; Cohen’s *d* = 1.30). Treatment with oxytocin did not affect adrenal mass (adrenal mass/body mass; all p’s>0.15) although we observed an effect of sex (F(1,58)=120.59; p<0.0001; partial η^2^ = 0.68; Cohen’s *d* = 2.83) with females having significantly larger relative adrenal mass compared to males.

**Table 1.** Mean (SEM) body mass at the start of injections (g), net body mass gain (g), and adrenal mass/body mass in male and female rats treated with oxytocin in PBS (OT 0.5 and 1.0 mg/kg) and in TRIOZAN^TM^ (OTT 0.5 and 1.0 mg/kg). Asterisks denote significant differences from control PBS males (p<0.05). ^a^ Denotes main effect of sex (p<0.001).

| Treatment | Sex | Sample Size | Initial body mass<sup>a</sup> (g) | Net body mass gain<sup>a</sup> (g) | Adrenal mass/body mass<sup>a</sup> |
| --- | --- | --- | --- | --- | --- |
| PBS | F | 8 | 252.13 (3.90) | 21.38 (3.01) | 0.025 (0.001) |
|  | M | 8 | 416.38 (13.49) | 55.00 (5.70) | 0.015 (0.001) |
| OT 0.5 | F | 8 | 248.38 (2.34) | 16.50 (2.01) | 0.028 (0.001) |
|  | M | 8 | 411.63 (11.25) | 37.88 (4.79)* | 0.013 (0.001) |
| OT 1 | F | 8 | 255.50 (2.06) | 11.50 (2.78) | 0.026 (0.002) |
|  | M | 8 | 406.00 (8.87) | 29.50 (6.64)* | 0.013 (0.001) |
| OTT 0.5 | F | 5 | 253.40 (5.85) | 10.80 (5.80) | 0.023 (0.003) |
|  | M | 5 | 390.20 (6.26) | 25.60 (5.60)* | 0.016 (0.001) |
| OTT 1 | F | 5 | 254.20 (6.03) | 14.80 (2.60) | 0.024 (0.003) |
|  | M | 5 | 393.80 (4.42) | 19.60 (9.50)* | 0.014 (0.001) |

### Oxytocin caused sedation

Injections of oxytocin caused sedation observed 5-10 mins post-injection (Uvnäs-Moberg et al., 1994). We calculated a sedation score using a binary score (0 = no sedation, 1 = sedation indicated by a clear lack of locomotor activity and rearing, eyes closed and rats appearing asleep) over the course of 10 days for a maximum possible score of 10. All the PBS control animals had a sedation score of 0 (i.e., no sedation). In contrast, all doses and delivery methods of oxytocin resulted in levels of sedation higher than 0 (Figure 1A). We therefore compared sedation scores for the oxytocin groups only. Male rats had higher levels of sedation than female rats for the low doses of oxytocin (0.5 mg/kg), regardless of delivery method (interaction between treatment and sex F(3,44)=4.08; p<0.05; partial η^2^ = 0.22; main effect of sex: F(1,44)= 50.04; p<0.0001; partial η^2^ = 0.53). Furthermore, for female rats oxytocin in TRIOZAN^TM^ (0.5 mg/kg) resulted in lower sedation relative to the same dose of oxytocin delivered in PBS (p=0.05; Cohen’s *d* = 0.88) and relative to the high dose of oxytocin in TRIOZAN^TM^ (1.0 mg/kg; p=0.0001; Cohen’s d = 2.78; Figure 1A).

**Figure 1.**
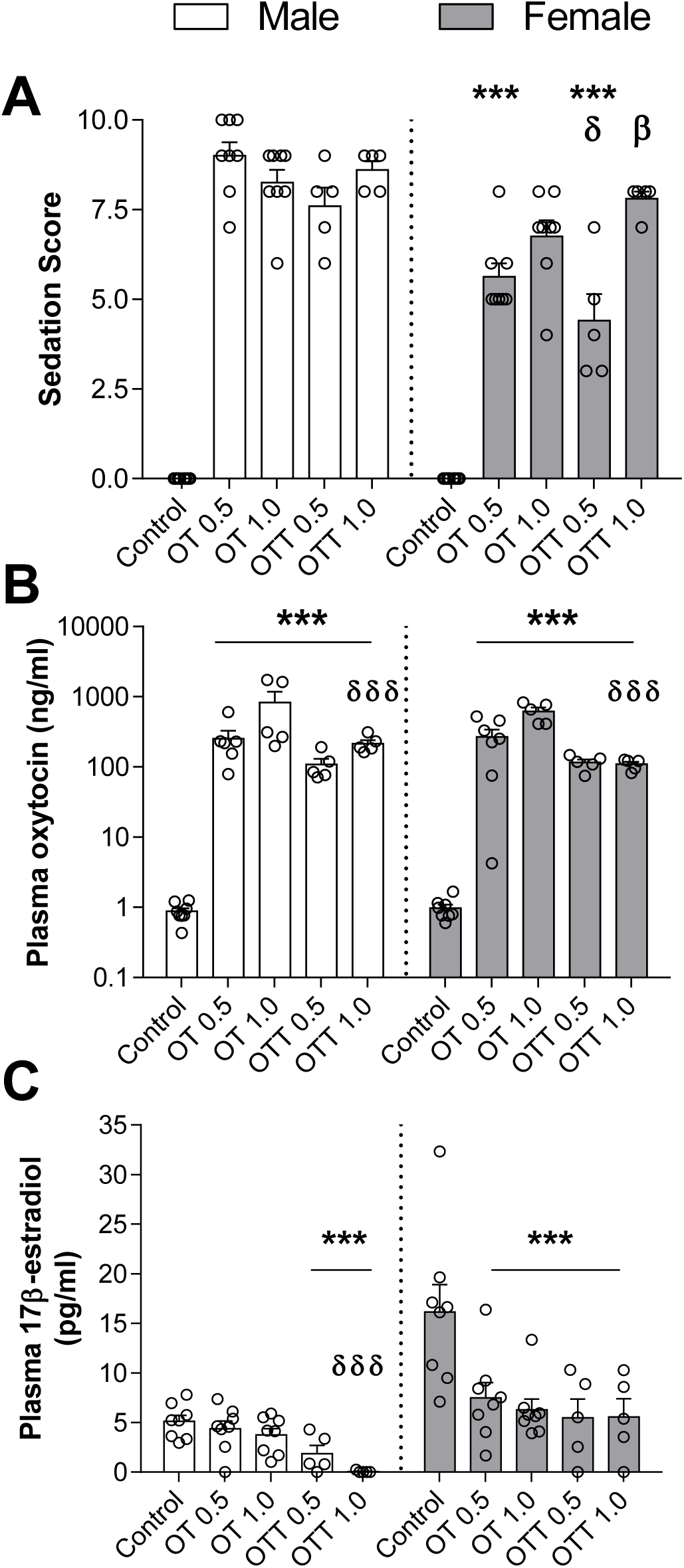
Oxytocin treatment caused sedation, increased plasma oxytocin and reduced plasma 17β-estradiol. **A.** Sedation scores in male and female rats immediately after injection with either PBS, oxytocin (OT, 0.5 or 1.0 mg/kg) or oxytocin in TRIOZAN^TM^ (OTT, 0.5 or 1.0mg/kg). Sedation scores higher than 0 (i.e. 0 = no sedation) were observed after oxytocin injection and not in the PBS controls. Males had higher sedation scores than females in the low doses of oxytocin, regardless of delivery method (*** indicates significantly different from males treated with oxytocin, OT and OTT 0.5 mg/kg; p<0.001). In females, oxytocin in TRIOZAN^TM^ (OTT 0.5 mg/kg) resulted in lower sedation scores relative to oxytocin in PBS (OT 0.5 mg/kg; δ indicates significant difference from OT 0.5; p=0.05) and relative to oxytocin in TRIOZAN^TM^ (1.0 mg/kg; β indicates significant difference between OTT 0.5 and OTT 1.0; p= 0.0001). **B.** Oxytocin treatment increased plasma levels of oxytocin in males and females relative to controls (*** p<0.001) and oxytocin in TRIOZAN^TM^ (OTT 1.0 mg/kg) resulted in lower plasma oxytocin compared to oxytocin in PBS (OT 1.0 mg/kg; δδδ indicates significant difference from OT 1.0; p<0.001). **C.** Oxytocin treatment reduced plasma 17β-estradiol levels in females and males relative to PBS controls (*** p<0.001). In females, all doses and delivery methods reduced 17β-estradiol levels. In males both doses of oxytocin in TRIOZAN^TM^ reduced 17β-estradiol in plasma. In addition, the highest dose of oxytocin in TRIOZAN^TM^ (OTT 1.0 mg/kg) resulted in lower plasma 17β-estradiol relative to oxytocin in PBS (OT 1.0 mg/kg; δδδ indicates significant difference from OT 1.0; p<0.001).

### Oxytocin injections increased oxytocin levels and decreased 17β-estradiol levels in both sexes

Daily oxytocin (both doses) injections increased circulating levels of plasma oxytocin in males and females regardless of delivery method (all p’s<0.001; main effect of treatment F(4,48)=358.3; p<0.0001; partial η^2^ = 0.97; Figure 1B). In males and females, both doses of oxytocin delivered in TRIOZAN^TM^ (0.5 and 1.0 mg/kg) resulted in lower plasma levels relative to oxytocin in PBS (p<0.01 and 0.0.001; Cohen’s *d* = 1.49 and 2.23, respectively). Oxytocin treatment (0.5 and 1.0 mg/kg in PBS and TRIOZAN^TM^) reduced plasma levels of 17β-estradiol in females relative to PBS controls, regardless of delivery method and estrous cycle phase (main effect of treatment F(4,22)=3.53, p<0.05; partial η^2^ = 0.39; Figure 1C). In males, oxytocin delivered in TRIOZAN^TM^ (0.5 and 1.0 mg/kg) reduced circulating levels of 17β-estradiol relative to PBS controls (p<0.05 and p<0.001, respectively; main effect of treatment F(4,29)=11.18, p<0.001; partial η^2^ = 0.61). The high dose of oxytocin delivered in TRIOZAN^TM^ (1.0 mg/kg) also reduced circulating levels of 17β-estradiol relative to oxytocin delivered in PBS (1.0 mg/kg; p<0.001).

### Oxytocin treatment increased brain levels of oxytocin when delivered in TRIOZAN^TM^ only

Oxytocin delivered in TRIOZAN^TM^ (1.0 mg/kg) increased oxytocin concentrations in the hypothalamus in males and females relative to oxytocin delivered in PBS (1.0 mg/kg; main effect of treatment F(4,49)=3.16, p=0.02, partial η^2^ =0.21; a significant covariate effect of 17β-estradiol (F(1,49)=5.84, p=0.02; partial η^2^ =0.10) to decrease oxytocin levels; Figure 2A). Oxytocin delivered in PBS did not affect the levels of oxytocin in the hypothalamus relative to the PBS controls in both sexes (p’s>0.3). To investigate how peripheral and hypothalamic levels of oxytocin compare between the treatment groups, we calculated the ratio of hypothalamus to plasma oxytocin content. Compared to the PBS controls, all oxytocin treatments significantly decreased the hypothalamus to plasma ratio in both males and females (all p’s<0.001; main effect of treatment F(4,36)=28.71, p<0.0001; partial η^2^ = 0.76; a significant covariate effect of 17β-estradiol (F(1,36)=4.11, p=0.05; partial η^2^ =0.10) to increase the hypothalamus to plasma ratio; Figure 2B). In addition, oxytocin in PBS (1.0 mg/kg) resulted in a lower hypothalamus to plasma ratio compared to the oxytocin delivered in TRIOZAN^TM^ (1mg/kg; p<0.01; Cohen’s *d* = 2.15). Using ELISA, we were not able to detect oxytocin levels in other homogenized brain regions (frontal region, hippocampus). To complement the total levels of oxytocin in the homogenized hypothalamus, we used immunofluorescence to investigate the effect of oxytocin treatments on the number of OT-ir cells in two hypothalamic nuclei, SON and PVN (Figure 2C and D). There was a weak correlation (r= −0.248; p=0.08) between plasma 17β-estradiol and the number of OT-ir cells in the PVN, therefore we used 17β-estradiol levels as a covariate in the analysis of OT-ir cells. In females, oxytocin in PBS and oxytocin delivered in TRIOZAN^TM^ did not affect the number of OT-ir cells in the SON and PVN (all p’s> 0.3; Figure 2C and D). In males, oxytocin in PBS (1.0 mg/kg; p=0.03; Cohen’s *d* = 2.00) and oxytocin delivered in TRIOZAN^TM^ (0.5 and 1.0 mg/kg; p= 0.02 and 0.03; Cohen’s *d* = 1.98 and 2.33, respectively) increased the number of OT-ir cells in the SON relative to PBS controls (main effect of treatment F(4, 19) =3.63; p< 0.05; partial η^2^ = 0.43; Figure 2D). Oxytocin delivered in TRIOZAN^TM^ (0.5 mg/kg) increased the number of OT-ir cells relative to oxytocin in PBS (0.5 mg/kg; p= 0.04; Cohen’s *d* = 1.56). In the PVN, oxytocin did not significantly affect the number of OT-ir cells in males (p’s>0.7).

**Figure 2.**
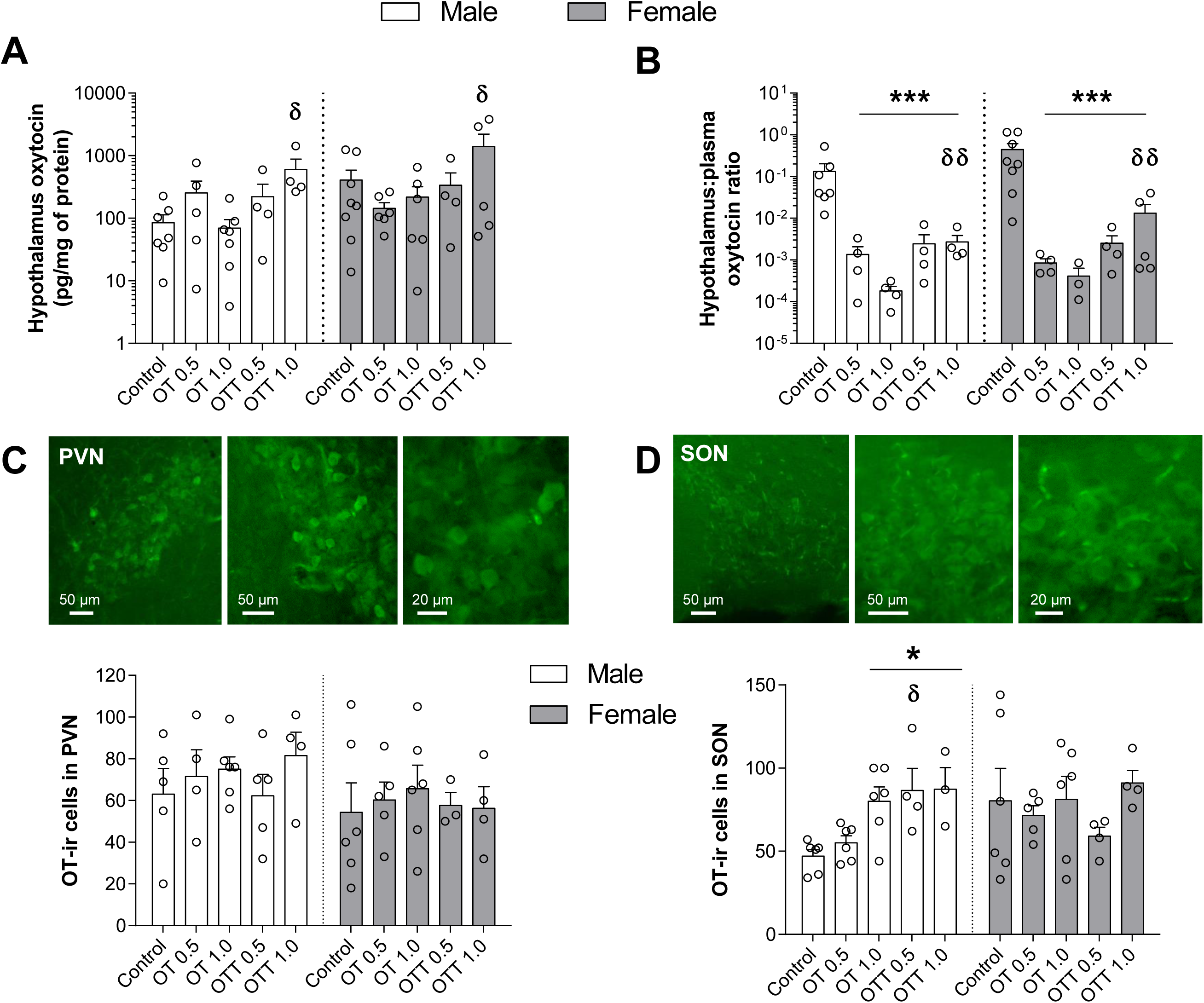
Oxytocin delivered in TRIOZAN^TM^ increased oxytocin levels in the hypothalamus. **A.** Oxytocin levels measured by ELISA in the hypothalamus were significantly higher in oxytocin in TRIOZAN^TM^ (OTT, 1.0mg/kg) compared to oxytocin in PBS in both sexes (OT, 1.0 mg/kg; indicated by δ, p<0.05). **B.** Ratio of hypothalamus to plasma levels of oxytocin were significantly lower in all oxytocin treatment groups (*** p<0.001) but oxytocin delivered in TRIOZAN^TM^ (OTT, 1.0 mg/kg) resulted in a higher ratio compared to oxytocin in PBS (OT, 1.0 mg/kg; indicated by δδ, p<0.01). **C.** In the PVN, the number of OT-ir cells was not affected by oxytocin treatment. **D.** There were more OT-ir cells in the SON of male rats treated with oxytocin in PBS (OT, 1.0 mg/kg) and in TRIOZAN^TM^ (OTT, 0.5 and 1.0 mg/kg) relative to PBS controls (p<0.05). The number of OT-ir cells in the SON was higher in oxytocin delivered in TRIOZAN^TM^ (OTT, 0.5 mg/kg) compared to oxytocin in PBS (OT, 0.5 mg/kg; indicated by δ, p<0.05). Representative images showing OT-ir cells in the PVN (**C**) and SON (**D**).

### Oxytocin treatment did not affect time spent in locomotion or grooming but decreased time spent rearing

Because we observed sedation shortly after oxytocin injection, we conducted social investigation tests 2 hours after oxytocin injection to ensure locomotion was not affected during the social investigation test. During the habituation period in the three-chambered arena, time spent in locomotion was not affected by oxytocin treatment (p’s>0.26; Table 2), but there was a trend for a main effect of sex indicating that females were more active than males (F(1,58)=3.53 p=0.06; partial η^2^ = 0.06). Oxytocin treatment did not affect the amount of time spent grooming (all p’s>0.41) but oxytocin in PBS (0.5 and 1.0 mg/kg) and in TRIOZAN^TM^ (1.0 mg/kg) decreased the time spent rearing during the habituation period relative to the PBS control (p’s=0.046, 0.018 and 0.014 respectively; Cohen’s *d* = 0.76, 1.34 and 1.34, respectively) in males and females (main effect of treatment F(4,58)=4.15; p<0.01; partial η^2^ = 0.22), and females reared more than males (main effect of sex F(1,58)=11.37, p<0.01; partial η^2^ = 0.16; for females a significant covariate effect of estrous cycle (F(1,27)=4.44, p=0.04; partial η^2^ =0.14)).

**Table 2.** Total (SEM) amount of time (in seconds) spent in locomotion, grooming, and rearing during the habituation of the three-chamber approach test in male and female rats treated with oxytocin in PBS (OT 0.5 and 1.0 mg/kg) and in TRIOZAN^TM^ (OTT 0.5 and 1.0 mg/kg). Asterisks denote significantly different from PBS (p<0.05). ^a^ Denotes main sex effect (p<0.01).

| Treatment | Sex | Sample Size | Locomotion (s) | Grooming (s) | Rearing<sup>a</sup> (s) |
| --- | --- | --- | --- | --- | --- |
| PBS | F | 8 | 202.24 (18.45) | 13.75 (2.66) | 169.80 (9.73) |
|  | M | 8 | 168.94 (13.41) | 13.97 (4.09) | 137.43 (16.70) |
| OT 0.5 | F | 8 | 185.90 (14.44) | 12.17 (3.13) | 141.74 (20.30)* |
|  | M | 8 | 171.14 (33.88) | 11.20 (3.76) | 85.18 (20.26)* |
| OT 1 | F | 8 | 184.48 (19.38) | 9.24 (2.67) | 119.18 (9.21)* |
|  | M | 8 | 129.49 (11.92) | 8.20 (2.11) | 81.97 (12.28)* |
| OTT 0.5 | F | 5 | 166.29 (21.60) | 14.95 (5.68) | 144.29 (15.33) |
|  | M | 5 | 157.94 (17.04) | 11.66 (2.73) | 116.69 (13.33) |
| OTT 1 | F | 5 | 149.51 (15.41) | 9.84 (7.28) | 112.86 (15.76) |
|  | M | 5 | 136.56 (22.45) | 22.41 (6.67) | 90.50 (16.86) |

### Oxytocin increased social investigation in males but not females

During habituation, the time spent in each of the three chambers was not significantly different between treatments indicating that animals did not have a preference for a particular side of the chamber (all p’s>0.4; data not shown). We quantified the time spent in the social stimulus third of the chamber and the time investigating the social stimulus rat (i.e. sniffing and touching the basket where the stimulus rat was confined) as two measures of social investigation. Regardless of treatment and sex, rats spent significantly more time in the chamber with the social stimulus rat compared to the empty or middle chambers (main effect of side F(2,116)=191.3, p<0.0001; partial η^2^ = 0.77; Table 3). Surprisingly, none of the oxytocin treatments affected the time spent in chamber with the social stimulus (p’s>0.13; Figure 3B). However, we did find effects of oxytocin on time spent investigating the stimulus basket. Oxytocin in PBS (0.5 mg/kg) increased the time spent investigating the stimulus basket in males (p=0.049; Cohen’s *d* = 1.83) but not female rats relative to PBS controls (p’s>0.6; main effect of treatment, F(4,58)=3.67, p<0.01, partial η^2^ = 0.20; interaction between treatment and sex, F(4,58)=2.50, p=0.05, partial η^2^ = 0.15; no significant main effect of sex, p>0.12; Figure 3C). Social investigation of the stimulus rat was not significantly different between oxytocin in PBS and oxytocin in TRIOZAN^TM^ groups (p’s>0.2).

**Figure 3.**
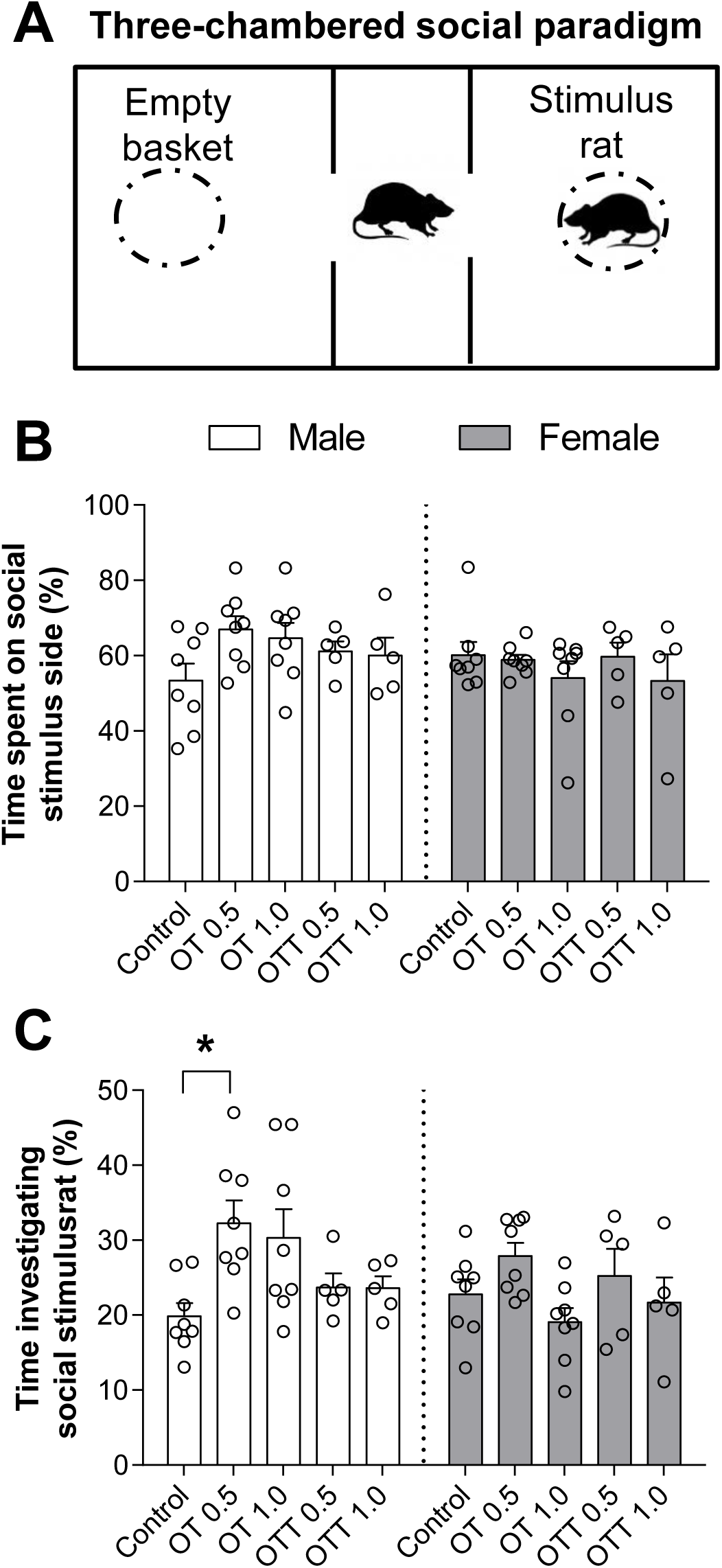
Oxytocin increased social investigation in male but not female rats in the three chambered social approach test. **A.** Three-chambered apparatus for quantifying social investigation. **B.** Oxytocin did not affect the percent time spent on social stimulus side. **C.** In males, oxytocin in PBS (OT 0.5 mg/kg) increased time sent investigating social stimulus relative to control (p<0.05).

**Table 3.** Percent time spent (SEM) in each of the chambers during the social behaviour test in male and female rats treated with oxytocin in PBS (OT 0.5 and 1.0 mg/kg) and in TRIOZAN^TM^ (OTT 0.5 and 1.0 mg/kg). Rats spent more time in the social stimulus side compared to the empty and middle chambers, regardless of treatment and sex (^a^ denotes a significant main effect of side; p<0.0001).

| Treatment | Sex | Sample Size | Time in Social Stimulus Chamber<sup>a</sup> | Time in Empty Chamber | Time in Middle Chamber |
| --- | --- | --- | --- | --- | --- |
| PBS | F | 8 | 60.13 (3.52) | 28.35 (3.03) | 11.52 (1.18) |
|  | M | 8 | 53.34 (4.50) | 23.79 (4.35) | 22.84 (5.82) |
| OT 0.5 | F | 8 | 58.83 (1.41) | 29.26 (1.91) | 11.9 (2.23) |
|  | M | 8 | 66.92 (3.51) | 22.17 (3.45) | 10.91 (1.98) |
| OT 1 | F | 8 | 54.00 (4.51) | 26.15 (3.01) | 19.84 (5.58) |
|  | M | 8 | 64.59 (4.13) | 19.21 (3.47) | 16.23 (3.41) |
| OTT 0.5 | F | 5 | 59.73 (3.70) | 27.92 (3.75) | 12.35 (2.47) |
|  | M | 5 | 61.12 (2.65) | 25.96 (2.10) | 12.92 (2.15) |
| OTT 1 | F | 5 | 53.30 (7.10) | 24.60 (5.20) | 22.09 (10.64) |
|  | M | 5 | 60.07 (4.72) | 27.36 (4.80) | 12.57 (2.79) |

### Oxytocin decreased hippocampal neurogenesis in males and females

The volume of the GCL was larger in males than in females (main effect of sex F(1,58)=18.01, p<0.0001, partial η^2^ =0.24; Table 4) and the volume of the ventral GCL was larger than the volume of the dorsal GCL (main effect of region; F(1,58)=212.5; p<0.0001, partial η^2^ =0.79). Oxytocin treatments had no effect on GCL volumes (p’s>0.11), therefore we present number of DCX+ cells. Oxytocin in PBS (1.0 mg/kg) decreased the number of DCX+ cells in the dorsal GCL relative to PBS control in males and females (p=0.015; Cohen’s *d* = 0.86; interaction between region and treatment F(4,54)=4.17, p<0.01, partial η^2^ = 0.24; Figure 4B). In the ventral region, oxytocin in PBS (0.5 and 1.0 mg/kg) and delivered in TRIOZAN^TM^ (1.0 mg/kg) decreased the number of DCX+ cells relative to PBS control in both sexes (p=0.0002, 0.0001, 0.002, respectively; Cohen’s *d*=0.95, 1.26, 0.72, respectively; Figure 4C). The latter effect of oxytocin in TRIOZAN^TM^ (1.0 mg/kg) relative to PBS was driven by the females (a priori comparisons; p<0.001, Cohen’s *d*= 1.59 in females; p=0.50 in males). Oxytocin in PBS (0.5 mg/kg) also decreased the number of DCX+ cells in the ventral region relative to oxytocin in TRIOZAN^TM^ (0.5 mg/kg) in males and females (p=0.002; Cohen’s *d*=0.85). Males had more DCX+ cells in the hippocampus than females (main effect of sex F(1,54)=9.98, p<0.01, partial η^2^ =0.16) and the ventral region had more DCX+ cells than the dorsal region in both sexes (main effect of region F(1,54)=385.9, p<0.0001, partial η^2^ =0.88). We also examined the maturity of DCX+ cells in the dorsal and ventral hippocampus (Table 4). Oxytocin (in PBS or TRIOZAN^TM^) had no effect on the percentage of DCX+ cells at each stage of maturity in males and females (p’s>0.4). There were significantly more type 3 DCX+ cells relative to type 1 and 2 (main effect of type of DCX+ cell; F(2,116)=23.09, p<0.0001; partial η^2^ =0.28).

**Figure 4.**
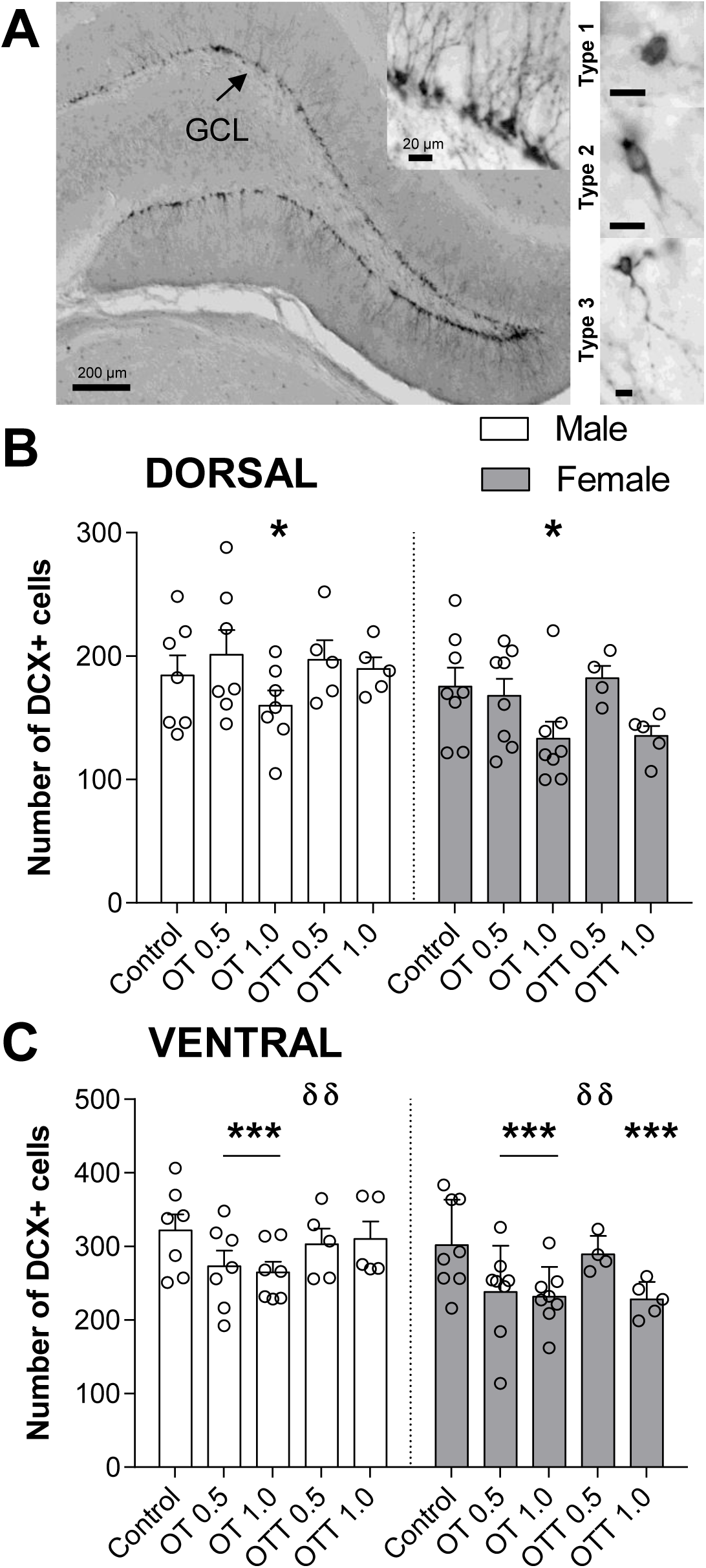
Oxytocin affected adult neurogenesis in a region- and delivery-dependent way. **A.** Representative images of DCX+ cells in the granule cell layer of the hippocampus and the three types of DCX+ cells based on morphology (scale bar is 10 μm). **B.** In the dorsal hippocampus, oxytocin in PBS (OT 1.0 mg/kg) decreased the number of DCX+ cells in males and females relative to PBS controls (p<0.05). **C.** In the ventral hippocampus, oxytocin in PBS (OT 0.5 and 1.0 mg/kg) decreased the number of DCX+ cells in males and females (p<0.001). In females, oxytocin delivered in TRIOZAN^TM^ (OTT 1.0 mg/kg) decreased the number of DCX+ cells relative to PBS controls (p<0.001). In both sexes, oxytocin delivered in TRIOZAN^TM^ (OTT 0.5 mg/kg) increased the number of DCX+ cells relative to oxytocin in PBS (OT 0.5 mg/kg; δδ indicates significantly different from OT 0.5, p<0.01).

**Table 4.** Mean (SEM) dorsal and ventral volumes (mm^3^) of the granule cell layer (GCL) of the right hippocampus and percent of DCX+ cells in different stages of maturity (proliferative, type 1; intermediate, type 2; post-mitotic, type 3) in male and female rats treated with oxytocin in PBS (OT 0.5 and 1.0 mg/kg) and in TRIOZAN^TM^ (OTT 0.5 and 1.0 mg/kg). Males had significantly larger GCL volume compared to females (main effect of sex; p<0.05). Ventral volumes were larger than dorsal volumes (main effect of region; p<0.0001). Oxytocin treatments did not affect the volume of the GCL (all p’s>0.11). There were significantly more type 3 DCX+ cells relative to type 1 and 2 (main effect of type of DCX+ cell; p<0.0001). Oxytocin treatments did not affect the proportion of DCX+ cells in each stage of maturation (all p’s>0.4).

| Treatment | Sex | Sample Size | Dorsal GCL Volume | Ventral GCL Volume | Dorsal |  |  | Ventral |  |  |
| --- | --- | --- | --- | --- | --- | --- | --- | --- | --- | --- |
|  |  |  |  |  | Type 1 | Type 2 | Type 3 | Type 1 | Type 2 | Type 3 |
| PBS | F | 8 | 0.58 (0.04) | 1.09 (0.07) | 22.50 (3.55) | 31.67 (1.92) | 45.83 (2.76) | 31.04 (2.78) | 35.00 (1.70) | 33.96 (2.11) |
|  | M | 8 | 0.71 (0.05) | 1.16 (0.11) | 24.22 (3.00) | 28.21 (2.71) | 47.57 (2.94) | 32.92 (3.18) | 33.33 (3.42) | 33.75 (2.69) |
| OT 0.5 | F | 8 | 0.63 (0.06) | 1.02 (0.08) | 24.72 (3.64) | 28.90 (2.75) | 46.38 (3.31) | 34.66 (2.98) | 32.56 (2.71) | 32.78 (2.70) |
|  | M | 8 | 0.65 (0.06) | 1.15 (0.08) | 26.08 (3.67) | 27.57 (1.75) | 46.36 (4.67) | 34.09 (3.59) | 33.47 (3.63) | 32.44 (2.17) |
| OT 1 | F | 8 | 0.52 (0.03) | 1.02 (0.06) | 30.07 (2.90) | 27.97 (2.55) | 41.96 (2.18) | 35.20 (2.70) | 34.17 (2.83) | 30.63 (2.58) |
|  | M | 8 | 0.68 (0.03) | 1.19 (0.10) | 26.88 (4.48) | 27.08 (2.15) | 46.04 (4.76) | 33.83 (3.22) | 34.65 (1.16) | 31.52 (3.16) |
| OTT 0.5 | F | 5 | 0.65 (0.10) | 1.09 (0.04) | 23.21 (6.57) | 29.06 (2.71) | 47.73 (5.48) | 30.43 (4.89) | 35.10 (3.12) | 34.47 (3.66) |
|  | M | 5 | 0.71 (0.06) | 1.23 (0.07) | 27.44 (6.53) | 24.74 (1.90) | 47.82 (5.03) | 32.33 (3.82) | 31.67 (2.47) | 36.00 (3.44) |
| OTT 1 | F | 5 | 0.61 (0.02) | 0.86 (0.05) | 27.08 (7.13) | 28.76 (3.26) | 44.16 (4.26) | 30.00 (5.53) | 35.00 (1.90) | 35.00 (4.94) |
|  | M | 5 | 0.64 (0.05) | 1.28 (0.08) | 27.33 (3.82) | 30.33 (1.43) | 42.33 (3.23) | 27.67 (4.67) | 32.67 (3.44) | 39.67 (2.32) |

### Hippocampal neurogenesis was associated with 17β-estradiol in females and social investigation in males

We found that increased numbers of DCX+ cells were significantly associated with increased 17β-estradiol in the dorsal (Pearson’s correlation; r(31)=0.44; p=0.01) and ventral (r(31)=0.48; p=0.005) hippocampus in female but not male rats (r(29)=-0.013, p=0.95; r(29)=-0.05, p=0.80, respectively; Figure 5A and B). After removing one outlier in females with high 17β-estradiol levels (∼32 pg/ml; Figure 5A and B include this outlier), the only correlation that remains significant is in the ventral region (r(30)=0.36, p=0.04). In females, the correlation between DCX+ cells and 17β-estradiol was observed in the PBS control (dorsal, r(6)=0.84, p=0.009; ventral, r(6)=0.70, P=0.05; although these were not significant with the female outlier removed: dorsal, r(5)=0.69, p=0.09; ventral, r(5)=0.67, p=0.10) but not in any of the oxytocin groups (p’s>0.4). We also found that increased DCX+ cells in the ventral hippocampus was associated with decreased social investigation in males (r(32)=-0.40, p=0.03) but not females (r(32)=0.07, p=0.71; Figure 5C). Finally, we found no significant correlations between neurogenesis (ventral or dorsal) and OT-ir cells in the PVN and SON.

**Figure 5.**
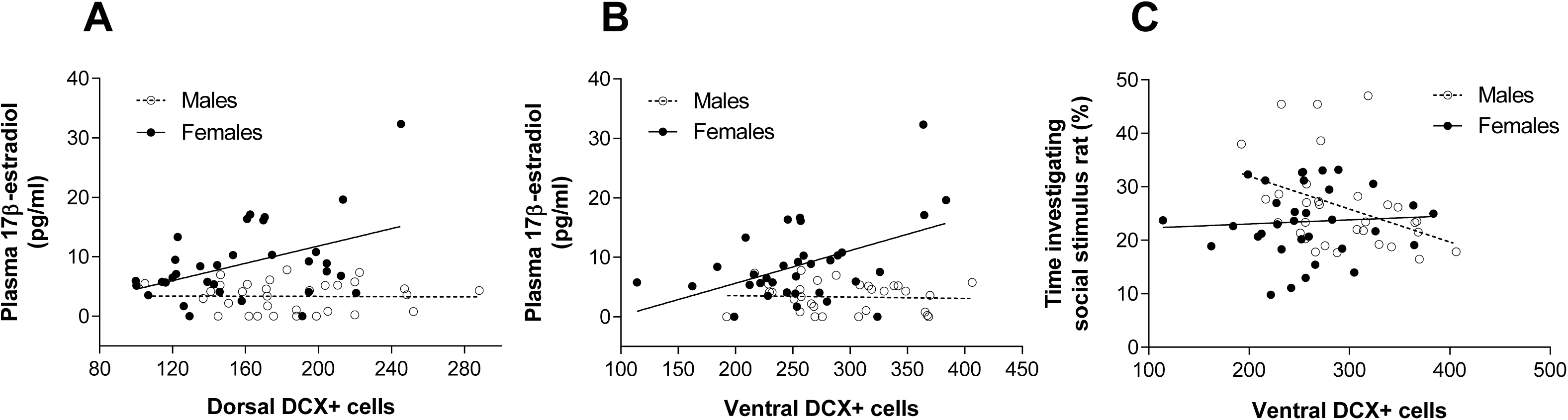
Number of DCX+ cells was significantly associated with plasma 17β-estradiol in females and social investigation in males. Increased numbers of DCX+ cells were significantly associated with increased 17β-estradiol in the dorsal (**A**, r(31)=0.44; p=0.01) and ventral (**B**, r(31)=0.48; p=0.005) in female but not male rats (p>0.80). In females, the correlation between DCX+ cells and 17β-estradiol was observed in the PBS control (dorsal, r(6)=0.84, p=0.009; ventral, r(6)=0.70, P=0.05) but not in any of the oxytocin groups (p’s>0.4). Increased DCX+ cell counts in the ventral hippocampus were associated with decreased social investigation in males (**C**, r(32)=-0.40, p=0.03) but not females (r(32)=0.07, p=0.71).

## Discussion

In the present study, we found that oxytocin treatment decreased body mass, increased social investigation and the number of OT-ir cells in the SON in male but not female rats. In contrast in both sexes, oxytocin reduced plasma 17β-estradiol levels and neurogenesis in the ventral hippocampus in a dose- and delivery-dependent way. Oxytocin treatment, regardless of delivery and sex, resulted in sedation effects immediately after injection. Oxytocin delivered in TRIOZAN^TM^ produced some effects that differed from oxytocin delivered in PBS. Principally, oxytocin delivered in TRIOZAN^TM^ reduced sedation and plasma oxytocin levels relative to oxytocin delivered in PBS. In addition, oxytocin in TRIOZAN^TM^ increased central measures such as hypothalamic oxytocin levels and ventral neurogenesis relative to oxytocin in PBS in males and females. Altogether, oxytocin resulted in sex-specific effects on social investigation and OT-ir cells in the SON but similar effects between the sexes on hippocampal neurogenesis, plasma 17β-estradiol and hypothalamus oxytocin levels, although some of these effects were dependent on the dose and delivery system used.

### Oxytocin increased social investigation in males but not females

In this study, both males and females displayed a preference for the social stimulus in line with previous research in rats (Lukas and Neumann, 2014) and mice (Crawley et al., 2007). We found that treatment with oxytocin for 10 days increased social investigation in male, but not in female, rats. The finding in males is consistent with previous research in male mice and rats (Lukas et al., 2011; Ramos et al., 2013). However, in our experiment, oxytocin did not increase social investigation in female rats, which is similar to the previous findings using central infusions of either oxytocin or an OTR antagonist (Lukas and Neumann, 2014). Thus, oxytocin appears to have no significant influence on social investigation in non-postpartum female rats, across a variety of paradigms and injection procedures. Consistent with our work, in rats, infusion of an OTR antagonist in the amygdala reduced social investigation in males only (Dumais et al., 2016a). These findings suggest that oxytocin manipulations influence social investigation in males but not females.

There are very few studies doing direct comparisons between males and females, nevertheless sex differences in the effects of oxytocin on neural activation have been observed in humans and rodents. In humans, intranasal oxytocin resulted in sex-specific effects on brain activity during reciprocated cooperation (Rilling et al., 2014). Oxytocin increased brain activity in males in areas important for reward, social bonding, arousal, and memory (the striatum, basal forebrain, insula, amygdala and hippocampus), whereas, oxytocin had no effect or decreased this response in certain areas in females (Rilling et al., 2014). In rats, oxytocin treatment also resulted in sex differences in neural activation patterns as males had more activation than females in the nucleus accumbens, insular cortex, pituitary, and some regions of the hypothalamus, whereas, females had more activation in the amygdala than males (Dumais et al., 2017). In addition, males have more OTR binding sites in hypothalamic and amygdala areas compared to females (DiBenedictis et al., 2017; Smith et al., 2017). In the current study, we found that oxytocin increased the number of OT-ir cells in the SON in males only. Oxytocin can autoregulate its own secretion in the hypothalamus by binding to OTR in male mice (Lopatina et al., 2010) and during parturition in female rats (Neumann et al., 1996). Therefore, it is possible that in male rats oxytocin treatment resulted in further oxytocin release in the SON via autoregulation but this positive feedback is not present in females (outside of parturition). Altogether, the lack of an effect of oxytocin in females on social investigation could be related to sex differences in the effects of oxytocin on OT-ir cells, OTR binding, autoregulation of oxytocin release, and neural activation.

### Oxytocin decreased ventral hippocampal neurogenesis in male and female rats

Contrary to previous studies in males, we found that oxytocin treatment decreased the number of immature neurons in the ventral hippocampus of both female and male rats. To our knowledge this is the first study investigating the effects of oxytocin on adult neurogenesis in female rodents. Previous studies in male rats have used similar doses and routes of exposure of oxytocin but different experimental designs (i.e., length of exposure, makers of neurogenesis, behavioural testing) making it difficult to directly compare our current results. In male rats, a single i.p. injection or central infusion of oxytocin increased cell proliferation in the ventral hippocampus (Leuner et al., 2012). Furthermore, oxytocin treatment for 7 days increased new neuron survival 1 or 3 weeks *after* the last injection (Leuner et al., 2012; Opendak et al., 2016). Similarly, oxytocin treatment for 14 days increased DCX+ cells in adult male rats that also underwent behaviour (Sánchez-Vidaña et al., 2016). Altogether, this suggests that in males, oxytocin increases neurogenesis after prolonged treatment (14 days) or 1 to 3 weeks after 7 days of treatment, however in the present study we found that oxytocin decreased neurogenesis (DCX+ cells) after 10 days of exposure. Thus, it is possible that in the current study we may have observed an increase after more prolonged exposure to oxytocin or a longer time prior to brain collection.

Interestingly, the effects of oxytocin on adult neurogenesis were mostly observed in the ventral hippocampus, similar to previous studies in rats (Leuner et al., 2012; Opendak et al., 2016). In contrast, other studies have seen effects in either both regions (in mice; Lin et al., 2017) or in the dorsal region only (in rats; Sánchez-Vidaña et al., 2016). The dorsal and ventral regions of the hippocampus have distinct functions (Bannerman et al., 2004; Fanselow and Dong, 2010; Hawley et al., 2012; Wu and Hen, 2014). Newly generated neurons in the dorsal and ventral hippocampus also project to different areas (reviewed in Strange et al., 2014). We observed a significant correlation between ventral neurogenesis and social investigation in males. It is possible that increased levels of ventral neurogenesis are associated with lower levels of social investigation in males. In addition, it is also possible that the regulation of ventral neurogenesis by oxytocin is related to other social-, stress-, or anxiety-related behaviours (Anacker et al., 2018; Kheirbek et al., 2013; Opendak et al., 2016; Snyder et al., 2011; Wu and Hen, 2014). As oxytocin also reduced 17β-estradiol levels (discussed below), and estradiol can regulate neurogenesis in both males and females (Barker and Galea, 2008; Eid et al., 2019; McClure et al., 2013; Ormerod et al., 2004), it is possible the effects we saw were related to oxytocin’s effects on circulating 17β-estradiol. However, in males this is unlikely as oxytocin reduced neurogenesis and 17β-estradiol either has no significant effect on neurogenesis in rats (after 15 days: Barker and Galea, 2008) or increases neurogenesis (after 5 days) in voles (Ormerod et al., 2004). Furthermore, no significant correlation was found between 17β-estradiol levels and DCX+ cell counts in males. In female rats, 17β-estradiol can suppress (after 15 days Barker and Galea, 2008) or increase neurogenesis (after 20 days, McClure et al., 2013; after 47 days, Eid et al., 2019). In the present study, we found a significant correlation between 17β-estradiol and DCX+ cell counts but this was mostly observed in the PBS controls. Although it is possible that in females the effects of oxytocin were mediated by effects on circulating 17β-estradiol, the lack of a correlation in the oxytocin groups suggest this was not a large contribution.

In the current study, we found no significant effects of oxytocin on the proportion of cells at different stages of maturation (proliferative, intermediate and mature) immediately after 10 days. In contrast, oxytocin treatment (1.0 mg/kg) for 14 days increased cell proliferation, number of mature DCX+ cells (i.e., cells with tertiary dendrites), and dendritic complexity in the dorsal hippocampus in male rats (Sánchez-Vidaña et al., 2016). In the latter study, animals were treated with oxytocin for a longer period (14 days vs 10 days present study) and were subjected to a battery of behavioural tests (including, forced swim test (FST)) which may have influenced the results. For example, behavioural despair tests (FST) can increase oxytocin levels in the brain of male rats (Yan et al., 2014). Altogether, our work suggests that oxytocin similarly regulates adult hippocampal neurogenesis in males and females but based on previous research, the effects may be affected by length of exposure, neurogenesis endpoints (proliferation, maturation or survival), and interaction with experiences.

### Nanoparticle increased oxytocin levels in the brain and reduced oxytocin sedation side effect

We found that delivery of oxytocin with the nanoparticle TRIOZAN^TM^ increased the levels of oxytocin in the hypothalamus and reduced sedation and plasma oxytocin levels relative to delivery with PBS. Thus, we saw oxytocin delivered in TRIOZAN^TM^ reduced peripheral effects by decreasing sedation and increased some central effects by increasing hypothalamic oxytocin levels relative to oxytocin delivered in PBS. Penetration of peripheral oxytocin into the brain is observed in small amounts (Neumann et al., 2013) calculated at ∼0.002-0.003% (Mens et al., 1983), 0.1% (Jones and Robinson, 1982), and 1.3% (Ermisch et al., 1985) but sufficient to induce behavioural and brain changes in hippocampal neurogenesis and brain activation (present study; Dumais et al., 2017; Leuner et al., 2012; Ring et al., 2006; Sánchez-Vidaña et al., 2016; Uvnäs-Moberg et al., 1992). Indeed, for a number of our measures we did not observe large differences between oxytocin delivered in PBS or in TRIOZAN^TM^.

In the current study, we observed sedation effects with both doses of oxytocin tested (0.5 and 1.0 mg/kg) similar to previous research showing that oxytocin causes sedation at high levels (0.25 and 1.0 mg/kg) and anxiolytic effects at lower levels (<0.004 mg/kg; Uvnäs-Moberg et al., 1994). However, female rats receiving oxytocin delivered in TRIOZAN^TM^ had less sedation compared to oxytocin in PBS which would suggest either a slow action or release of oxytocin in TRIOZAN^TM^ or a peripheral effect of oxytocin to cause sedation. There is evidence that oxytocin regulates skeletal muscle regeneration (Elabd et al., 2014) and that sleep is regulated by peripheral tissues such as skeletal muscle in mice (Ehlen et al., 2017). The nanoparticle TRIOZAN^TM^ can protect molecules from degradation (Ovensa Inc., 2018) and plasma levels of oxytocin were lower in the oxytocin in TRIOZAN^TM^ groups compared to oxytocin in PBS, which may suggest that oxytocin is more actively transported to tissues including the brain where it can either be stored or slowly released.

In our study, one limitation is that we cannot distinguish between endogenous and exogenous oxytocin. One possibility is that oxytocin delivered in TRIOZAN^TM^ increased endogenous production in the hypothalamus via indirect mechanisms (e.g., through projections from the periphery to the brain). Altogether, our work suggests that oxytocin in TRIOZAN^TM^ penetrated the brain but resulted in different behavioural and brain outcomes compared to oxytocin in PBS.

### Interactions between oxytocin and 17β-estradiol

In the present study, oxytocin treatment decreased plasma levels of 17β-estradiol in both sexes. There is evidence that oxytocin and estrogens interact (reviewed in Engel et al., 2019) but to date, research has focused on the effect of estrogens on the oxytocin system (Lim and Young, 2006; McCarthy et al., 1996; Tribollet et al., 1990). In human females, oxytocin levels decreased during times of lower estradiol phases during the menstrual cycle (meta-analysis by Engel et al., 2019) and with menopause (Maestrini et al., 2018). To our knowledge, our study is the first demonstration that oxytocin regulates circulating 17β-estradiol levels indicating that the interaction between these hormones is bidirectional. It is possible that the oxytocin regulation of neurogenesis is affected by endogenous levels of estradiol and potentially other sex steroids, which should be investigated.

## Summary

We found sex differences in the oxytocin regulation of social investigation and OT-ir cells in the SON of the hypothalamus. Oxytocin increased social investigation and the number of OT-ir cells in the SON in males but not females. In contrast, oxytocin decreased plasma 17β-estradiol and neurogenesis in the hippocampus of both males and females. We also show that the nanoparticle TRIOZAN^TM^ increased oxytocin levels in the hypothalamus in males and females and reduced sedation in females indicating that it can be a viable delivery system for oxytocin. Our study contributes to the evidence that oxytocin does not promote social investigation in both sexes.

## Acknowledgements

This study was funded by a Natural Sciences and Engineering Council of Canada grant to LAMG (RGPIN 203596-13). We thank Ovensa Inc. for the generous gift of TRIOZAN^TM^, Samantha Baglot and Dr. Steve Wainwright for helpful discussions on the design and results of the study, and Elizabeth Perez and Rebecca Rechlin for help with microscopy.

